# Comprehensive evolutionary analysis of complete Epstein Barr virus genomes from Argentina and other geographies

**DOI:** 10.1101/2021.03.05.434158

**Authors:** Ana Catalina Blazquez, Ariel José Berenstein, Carolina Torres, Agustín Izquierdo, Carol Lezama, Guillermo Moscatelli, Elena De Matteo, Mario Alejandro Lorenzetti, María Victoria Preciado

**Author notes:** Joint first authorship (Ana Catalina Blazquez, Ariel José Berenstein). Joint last authorship (Mario Alejandro Lorenzetti, María Victoria Preciado). Corresponding author: Maria Victoria Preciado, Instituto Multidisciplinario de Investigaciones en Patologías Pediátricas (IMIPP), CONICET-GCBA, Laboratorio de Biología Molecular, División Patología, Hospital de Niños Ricardo Gutiérrez. Gallo 1330, C1425EFD. Buenos Aires, Argentina.

## Abstract

Epstein Barr virus sequence variability has been deeply studied throughout the past years in isolates from various geographic regions and consequent geographic variation at both genetic and genomic levels has been described. However, isolates from South America have been underrepresented in these studies. Here, we sequenced 15 complete EBV genomes that we analyzed, by means of a custom-built bioinformatic pipeline, together with publicly available raw NGS data for 199 EBV isolates from other parts of the globe. Phylogenetic relations of the genomes, geographic structure and variability of the data set, and evolution rates for the whole genome and each gene were assessed. The present study contributes to overcome the scarcity of EBV complete genomes from South America, and hence, achieves the most comprehensive geography-related variability study by determining the actual contribution of each EBV gene to the geographic segregation of the entire genomes. Moreover, to the best of our knowledge, we established for the first time the evolution rate for the entire EBV genome, on a host-virus codivergence-independent assumption, and statistically demonstrate that evolution rates, on a gene-by-gene basis, are related to the encoded protein function. Considering evolution of dsDNA viruses with a codivergence-independent approach, may lay the basis for future research on EBV evolution. Additionally, this work also expands the sampling-time lapse of available complete genomes derived from different EBV-related conditions, a matter that until today, prevents for detailed phylogeographic analysis.

## Introduction

*Human gammaherpesvirus 4*, commonly referred to as Epstein-Barr virus (EBV), is the type species member of the *Lymphocryptovirus* genus, within the *Herpesviridae* family. After primary infection, EBV establishes life-long latency in memory B-lymphocytes in the human host and is present in over 90% of the world’s population; a fact that has rendered EBV as one of the most successful human viruses, which is assumed to be co-evolving with its host since the origins of mankind (Abdullah et al. 2017). In developing regions primary infection often occurs during early childhood and is not usually associated with clinical symptoms, although mild cases of infectious mononucleosis (IM) may occur. On the other hand, in developed regions, where primary infection is usually delayed until adolescence or early childhood, severe cases of IM are more frequent (Luzuriaga and Sullivan 2010). Although latency does not represent a significant risk in immunocompetent individuals, a co-factor involvement has been suggested in the development of neoplastic pathologies, such as Burkitt’s lymphoma (BL), Hodgkin’s lymphoma (HL), posttransplant lymphoproliferative disorder (PTLD) and cancers of epithelial origin including nasopharyngeal carcinoma (NPC) and some cases of gastric carcinomas (Farrell 2019).

Based on sequence variations located in EBNA2 gene and the EBNA3 gene family, EBV is broadly classified in EBV1 and EBV2. Further research at the genetic level described an association of specific variants with an augmented occurrence in tumoral samples, hence suggesting the possibility of an increased oncogenic potential (Xu et al. 2019; Hui et al. 2019). However, the role of EBV in the etiology and progression of these malignant processes is still not fully understood. Prevalence of EBV-associated malignancies, the percentage of EBV associated cases and viral variants vary between geographical regions worldwide; a fact that suggests the possibility of neoplastic-related EBV variants in different geographical regions, perhaps acting in synergy with genetic and/or environmental factors (Telford et al. 2020).

In past years, many studies from different parts of the world addressed the analysis of specific signature genes in the quest for viral variants strongly linked to different malignant pathologies (M. A. Lorenzetti et al. 2014; Neves et al. 2017; Chang et al. 2009). However, identifying EBV variants in tumor samples requires a better understanding of the viruses’ natural variation in the context of the entire genome so that potentially significant mutations can be distinguished from natural variation in the virus genome.

In recent years, with the advent of next generation sequencing (NGS) methods, it’s been possible to sequence the entire EBV genome and, to date, more than 1000 complete genomic sequences isolated from different pathologies and geographical regions are available in GenBank. However, most of these sequences come from Asian isolates, while South American isolates are deeply underrepresented, since they merely account for an approximate 2% of EBV complete genomic sequences available in the GenBank database. Hence, we expended the available data on EBV from Argentina and performed a comprehensive evolutionary analysis of the virus from several regions of the world, which allowed us to study its evolution history on a complete genome and on a gene-by-gene basis.

## Results

### 1. A call for a unified raw data sequence analysis

For the first time in our country, fifteen EBV-associated samples from pediatric Argentinean subjects were sequenced by NGS methods and were further analyzed with an automated and customized pipeline for viral data. To compare the variability of our genomes in the context of EBV sequences from other geographies, 199 publicly available raw-NGS data were downloaded from SRA-NCBI database and re-analyzed together with our raw-data making use of our new custom EBV-bioinformatics pipeline (see methods). Using this procedure, a total of 189 samples were classified as having type 1 EBNA2 and EBNA3s genes, 21 were type 2 EBNA2 and EBNA3s and only 4 were classified as recombinant genomes with type 1 EBNA2 and type2 EBNA3s. Regarding the 15 new Argentine sequences, 9 were classified as EBV1 and 6 were EBV2 (Supplemental Fig. 1).

Following viral typing, consensus sequences were constructed and aligned. Two different settings were explored: creating a multiple sequence alignment (MSA) with direct consensus sequences downloaded from GenBank (GB) and a MSA constructed with all sequences re-analyzed with a single pipeline (SP). Interestingly, SP sequences uniformly analyzed from fastq files produced a more reliable alignment than directly aligned consensus sequences from GenBank. In the first place, GB consensus sequences produced a longer alignment due to multiple gap insertions, which compensate for methodological discrepancies that arise when different pipelines are used (Fig. 1A and B). Moreover, while SP alignment preserves more than 82% of its positions perfectly aligned (zero entropy), GB consensus keeps only a 28% of its extent perfectly aligned (Fisher Exact Test; OR= 11.8 IC_95%_ [11.3 - 11.7], *P*-value <2.2 10^-16^). Concerning variability and SNPs, SP alignment reports 30824 SNPs, while the GB consensus alignment reports 169845 SNPs, most of which were located in repetitive regions; this made the GB alignment far more variable (Fisher Exact Test: OR= 0.086 IC95% [0.085 - 0.088], *P*-value <2.210^-16^). The latter case lacks biological plausibility, since the double strand DNA genome and the known stability of EBV is not compatible with the idea of having 70% of variable regions along its genome. On the other hand, a major lack of homogeneity was observed with the GB consensus procedure which denotes a significant number of entropic positions and a right-shifted entropy distribution in comparison with SP alignment (Fig. 1C). When only analyzing variable positions in both alignments (Entropy >0), significantly differences were found between distributions (Wilcoxon test, p-value< 2.2e-16) (Fig. 1D). All together, these results point out the drawbacks that arise when GenBank consensus sequences are directly considered to build a MSA for subsequent analysis, and call for the need of an unified process starting from raw fastq files.

**Figure 1.**
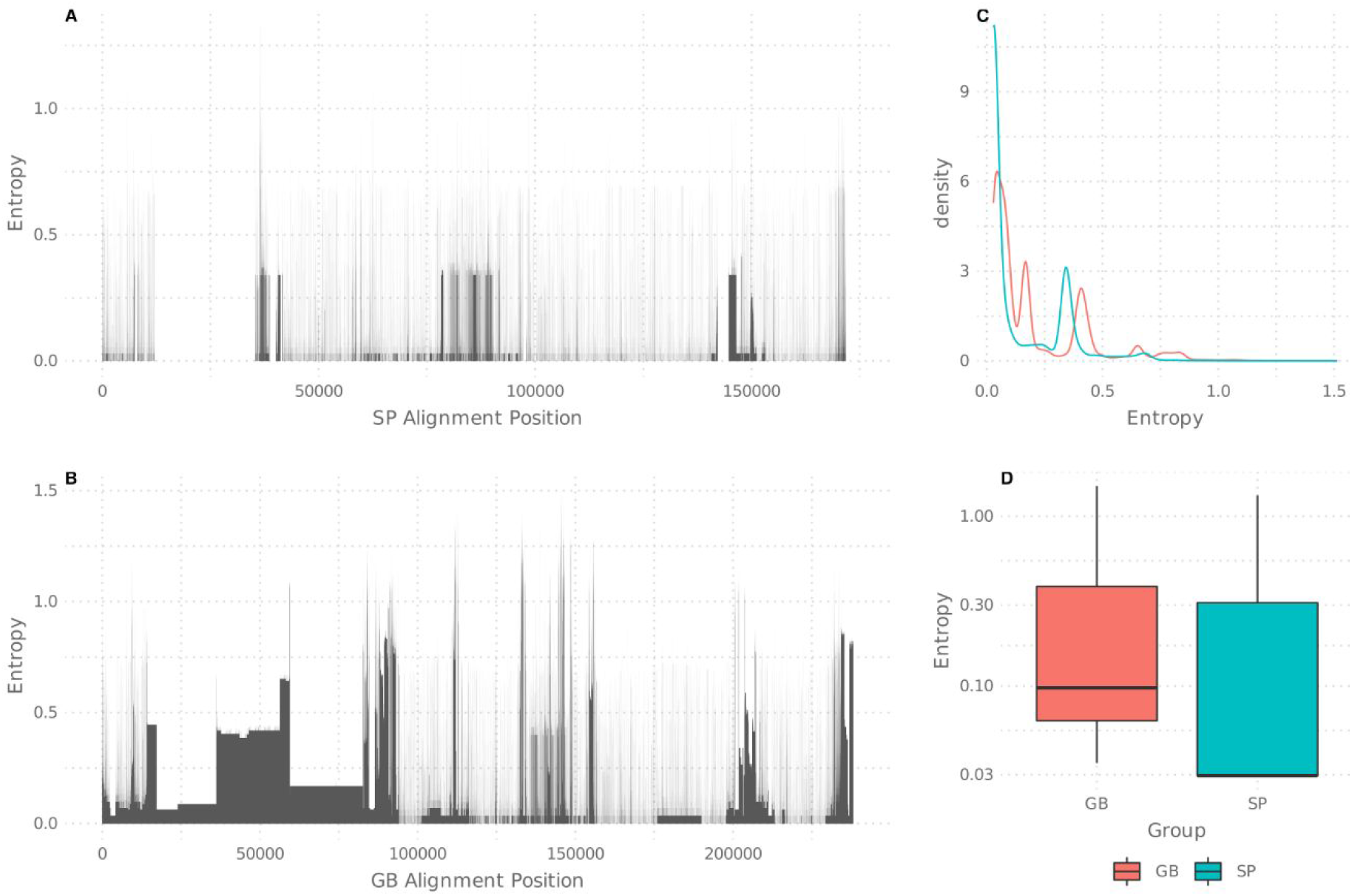
Alignment entropy analysis. (*A*) Entropic positions in the single pipeline (SP) generated alignment. (*B*) Entropic positions in the alignment directly generated with GenBank (GB) downloaded sequences. Highly entropic positions are depicted as black spikes or blocks. (*C*) Entropy density chart for both, SP and GB alignments considering all alignment positions. (*D*) Entropy comparison only considering positions with entropy >0, depicted in box plots.

### 2. Phylogenetic relations, evolution and geographic segregation

Following the construction of the all-new consensus sequences, and removal of recombinant genomes, the phylogenetic relationship between the Argentinean and worldwide genomes was inferred. As expected, the phylogenetic tree depicted two major clusters, EBV1 and EBV2. Within the EBV1 clade, two major supported subclusters were observed, with some inner groups with defined geographic structure; the first subcluster was composed mainly by Asian sequences and the second subcluster was more cosmopolitan, including intermingled sequences from Europe, Australia and the Americas-including some sequences from Argentina- and a supported inner group of African sequences. Furthermore, the first subcluster showed four subgroups: a small cluster formed by sequences from Western Asia (India, Saudi Arabia), two larger groups representing Eastern Asia (China, Japan, South Korea, Taiwan and Hong Kong) and Southeast Asia (Indonesia, Papua New Guinea and Singapore), and a group of sequences from Europe and Australia. In addition, three sequences from Argentina associated with viruses from Southeast Asia (Fig. 2). Regarding the EBV2 clade, most of the sequences were from Africa and Argentina, with minor representation of other geographic regions; however, the scarce number of EBV2 genomes prevented further statistical confidence, hence subsequent geographic analysis was performed considering EBV1 genomes only.

**Figure 2.**
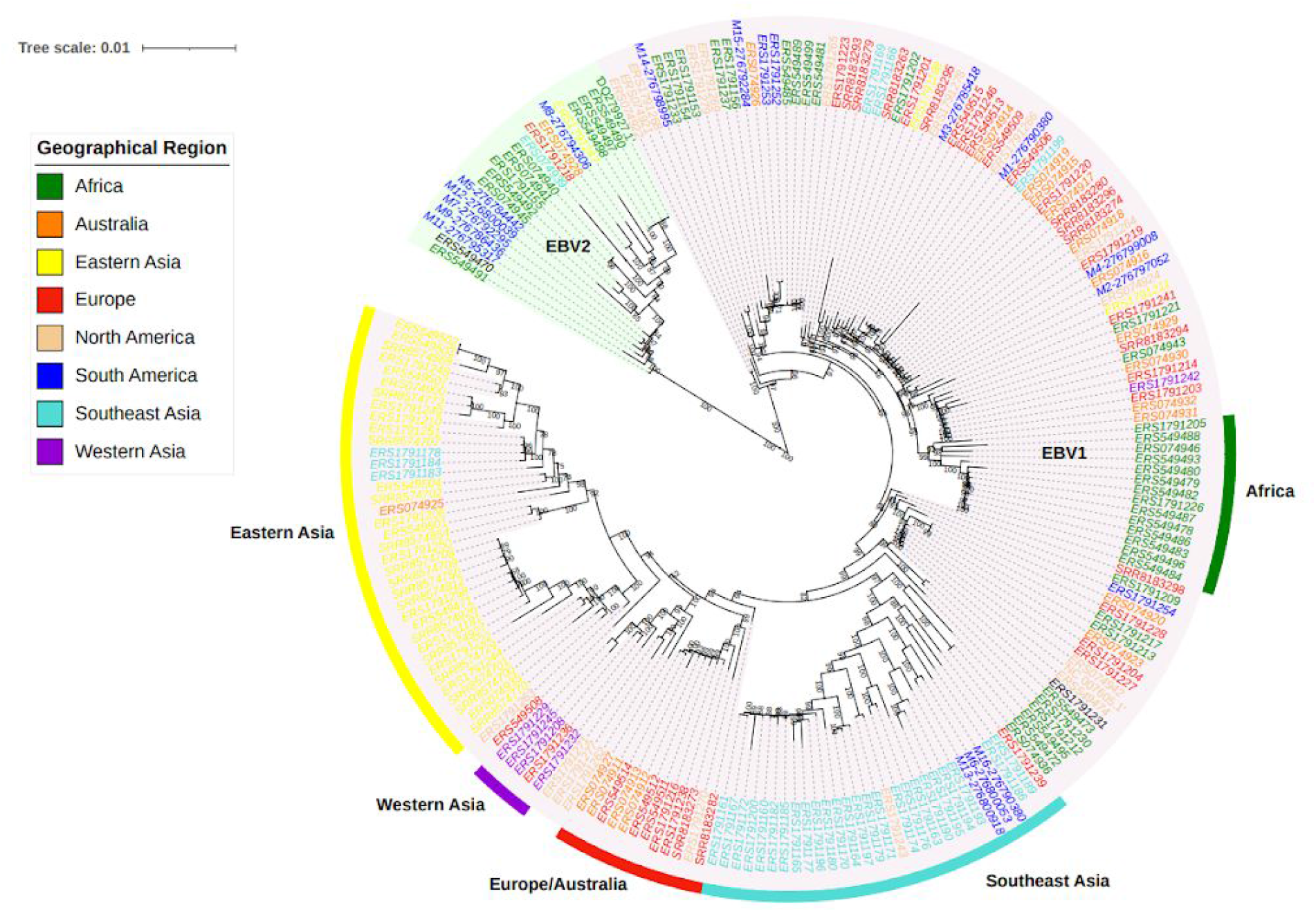
Phylogenetic reconstruction of 214 complete EBV genomes from different geographies. Phylogenetic tree constructed under Maximum likelihood method and 1000 ultrafast bootstrap resampling. Only values over 70 are shown. The green shaded clade contains the EBV-2 sequences and the pink shaded clade is EBV1. The color of each sequence represents the geographical region of the viral isolate. One sequence without a reliable origin of isolation is labeled in black. The five sub-clades were highlighted using different external colored bars. South American sequences correspond to isolates sequenced in the present study or were previously sequenced isolates from Argentina.

In order to statistically assess the number of variants within each region, variants were obtained from the VCF file by mapping the reads against the EBV1 reference sequence and then geographically stratified. This geographical variability was statistically significant, where sequences from Africa, Europe, Australia, North America and South America, presented a similar amount of variants when compared to the reference, but on the other hand, Asian genomes (South East Asia, Eastern Asia and Western Asia) contained significantly more variants when compared to the reference genome (Kruskal-Wallis test, *P*<2.2e-16) while presented the lesser intra-group variability, as denoted by the height of the box plot (Fig. 3A) (Supplemental Table 1).

**Figure 3.**
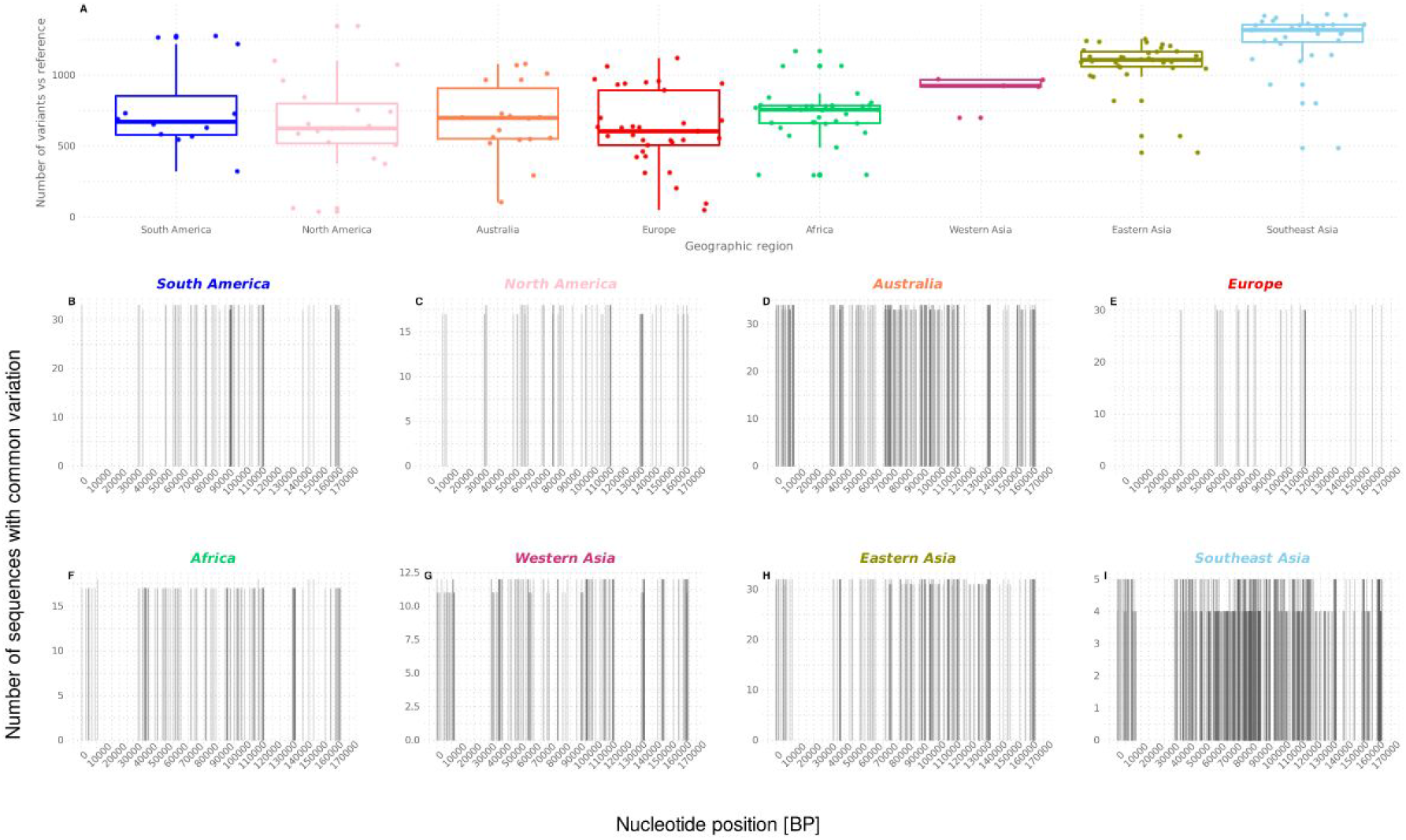
Quantification of EBV1 genomic variation regarding geographic origin. (*A*) Quantification and comparison of the amount of variants from different regions. Results are depicted in box plots. (*B-I*) Positioning of common variants against EBV1 reference along the genome for each geographical region.

To further elucidate the positioning of common variants within the EBV1 genome for each geographical region, independent alignments of geographically tagged sequences were generated and compared to the reference sequence using the custom built Consensus Compare tool (Fig. 3B-I). As observed, sequence variation is not homogeneously distributed along the genome in the different geographies. For instance, Asian sequences presented the most variable pattern of common variation along the genome; namely, the region between 120000pb-140000pb, which codes for late lytic genes (BDLF2, BDLF1, BcLF1, BcRF1, BTRF1, BXLF2, BXLF1, BVRF1, BVLF1, BVLF2, BdRF1 and BILF2), depicted variants in the South Asian, Eastern Asian and Western Asian sequences that were not present in the other geographical groups.

As a complementary analysis to the phylogenetic tree, in order to further assess the geographic structure in the EBV1 group, a PCA analysis was conducted. As shown in Figure 4A and 4B, PC1 (20.08%) mainly segregated the Asian genomes from the rest of the world’s sequences and PC2 (9.95%) allowed for the further discrimination of the Asian group into the three previously described subgroups, furthermore, both PC1 and PC2 distributions differ significantly among geographies (Kruskal-Wallis test, *P*-value: 8.58e-22 and 5.88e-17 respectively). Following this result, a pairwise Wilcoxon test was performed among all geographic regions. Such analysis quantified the observation that Eastern and Western Asian, Southeast Asian, as well as cosmopolitan genomes differ significantly on their first and second PCA distributions (Fig. 4C and 4D, Supplemental Table 2). According to the observed results, which supported the differential segregation of Asiatic EBV genomes from those from other continents, a Discriminant Analysis of Principal Components (DAPC) was performed with the aim of identifying the regions of the genome that drive the genetic divergence between both groups (Jombart, Devillard, and Balloux 2010). The number of PCs retained was defined according to the proportion of successful reassignment corrected for the number of retained PCs (alpha score); such a procedure was conducted for both, to improve the discrimination of the groups and to avoid overfitting. According to this criteria only one DAPC (DAPC1) was retained, which was sufficient to summarize the gene diversity between the Asian continent and the rest of the geographical regions. The probability of allocation of the Asian and non-Asian genomes was 84.5% and 89.65%, respectively. A ROC curve was constructed to estimate the potential of this classifier system, obtaining an AUC of 0.93 (0.88-0.97). (Fig. 5A). After that, structural SNPs were identified by means of “snpzip function of Adegenet package” (Jombart, Devillard, and Balloux 2010) which allowed for the identification of the four informative coding regions that were relevant in the segregation of Asian and cosmopolitan non-Asian clades (Fig. 5B). The first region was located between 43,281 to 60,240 bp with respect to the reference genome and variants located in this region were mainly concentrated in the BPLF1 gene. The second region was located between 95,719 to 97,415 bp and variants in this region were concentrated in EBNA-1 gene. The third group was located between 117,949 to 156,739 bp; although a large number of lytic genes are found in this region, variants mainly concentrated to BDLF3, BcRF1 and BXRF1 genes. Finally, the fourth region was located in the terminal region of the genome (168,236 to the end with respect to the reference), where the genes that encode for the oncogenic proteins LMP1, LMP2A and LMP2B are located. Additionally, isolated variants which also contributed to this differentiation were found in other regions of the genome with a lesser frequency (Supplemental Table 3).

**Figure 4.**
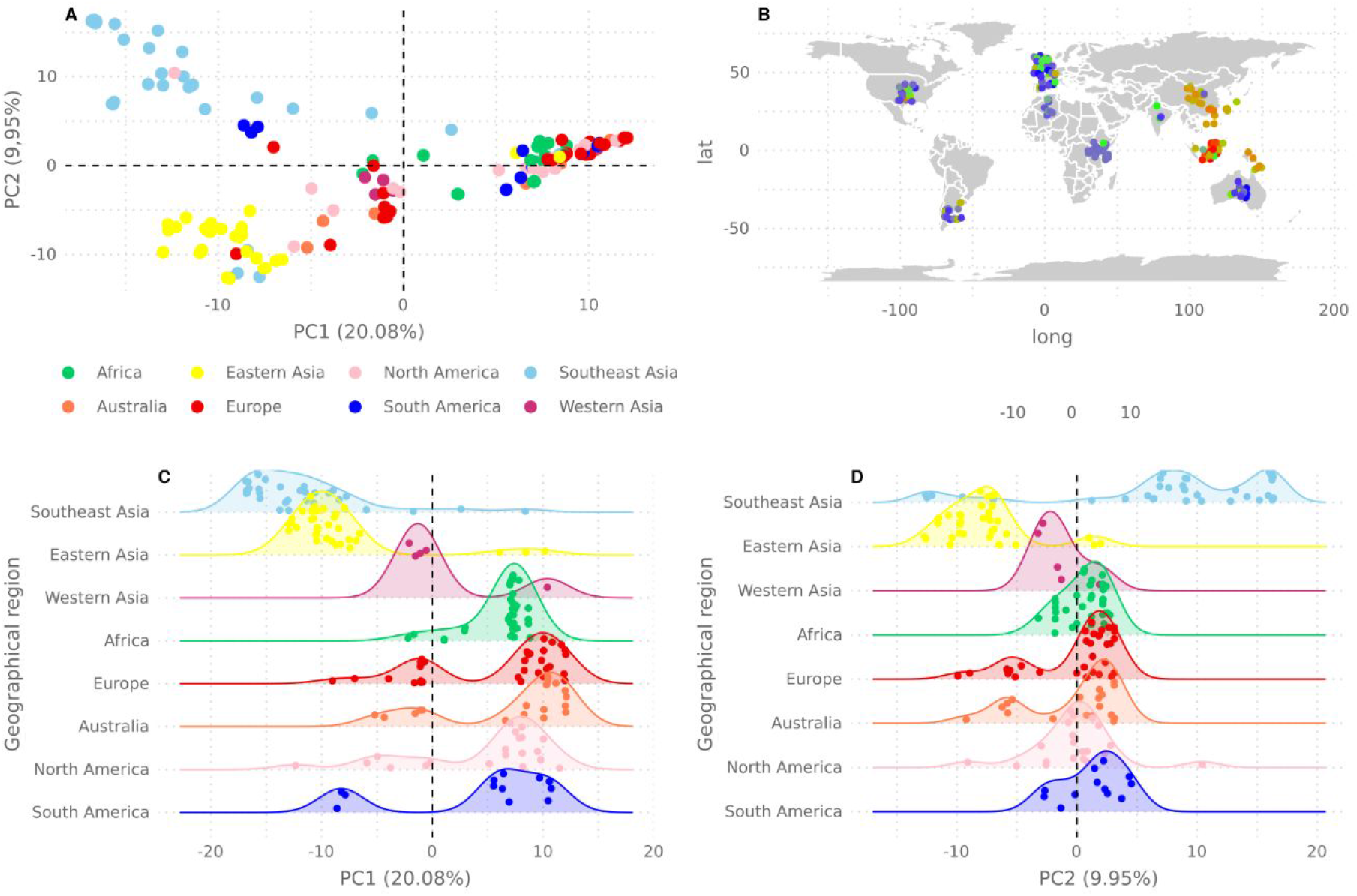
Principal component analysis. (*A*) PCA of EBV1 sequences showing PC1 and PC2. Sequences are coloured according to their geographic origin. (*B*) Geographical mapping the segregational potential of PC1 and PC2. (*C-D*) Quantification of the differences in PCA distribution, (*C*) for PC1 and (*D*) for PC2.

**Figure 5.**
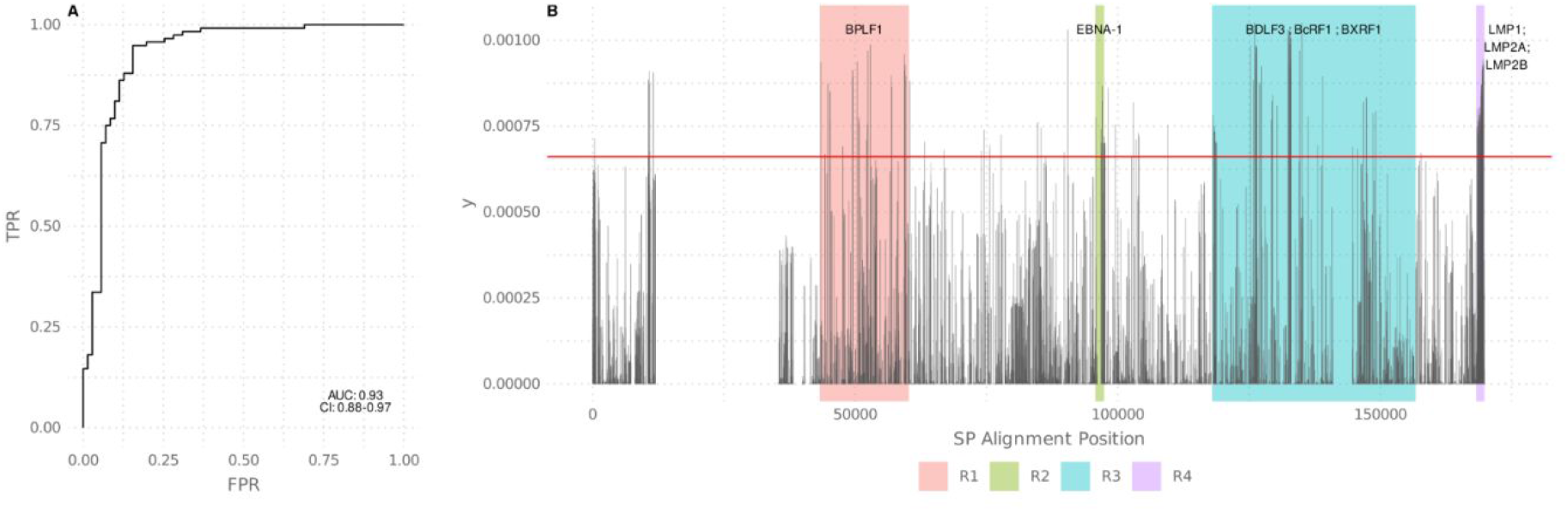
Discrimination analysis of principal components (DAPC). (*A*) ROC curve depicts the segregation potential of the DAPC main component between Asian and non Asian EBV1 sequences. *(B) DAPC* Variance contribution of each site (SNP) along the EBV1 genome. The red horizontal line indicates the threshold (0.00066) above which structural SNPs were identified. Colored boxes delimit the 4 informative coding regions of the genome. Most relevant genes within each region are indicated on the top of each box.

### 3. Genomic evolutionary rates

Given that each gene contributed differently to geographic segregation, we sought to study the evolutionary history of the entire genome and of each individual gene. For this purpose, the evolution rate for each individual gene and for the entire genome were calculated (Fig. 6A, Supplemental Table 4). In total 40 of the 60 analyzed genes exhibited evolutionary rates above the calculated mean for the entire genome (mean: 9.09e-6 s/s/y, 95% HPD interval: [4.08e-6, 1.56e-5]). In particular, BZLF1, BDLF3 and BcRF1 were amongst those genes with the highest evolutionary rates; interestingly, most of them are involved in viral immune escape as reported in the GO database. On the other hand, a total of 20 genes exhibited a similar evolutionary rate than that of the entire genome, with BFLF1, BMRF1 and BNRF1 being amongst those with the lowest evolutionary rates. As a whole, these genes are mainly involved in viral processes such DNA replication and the assembly of the viral particle.

**Figure 6.**
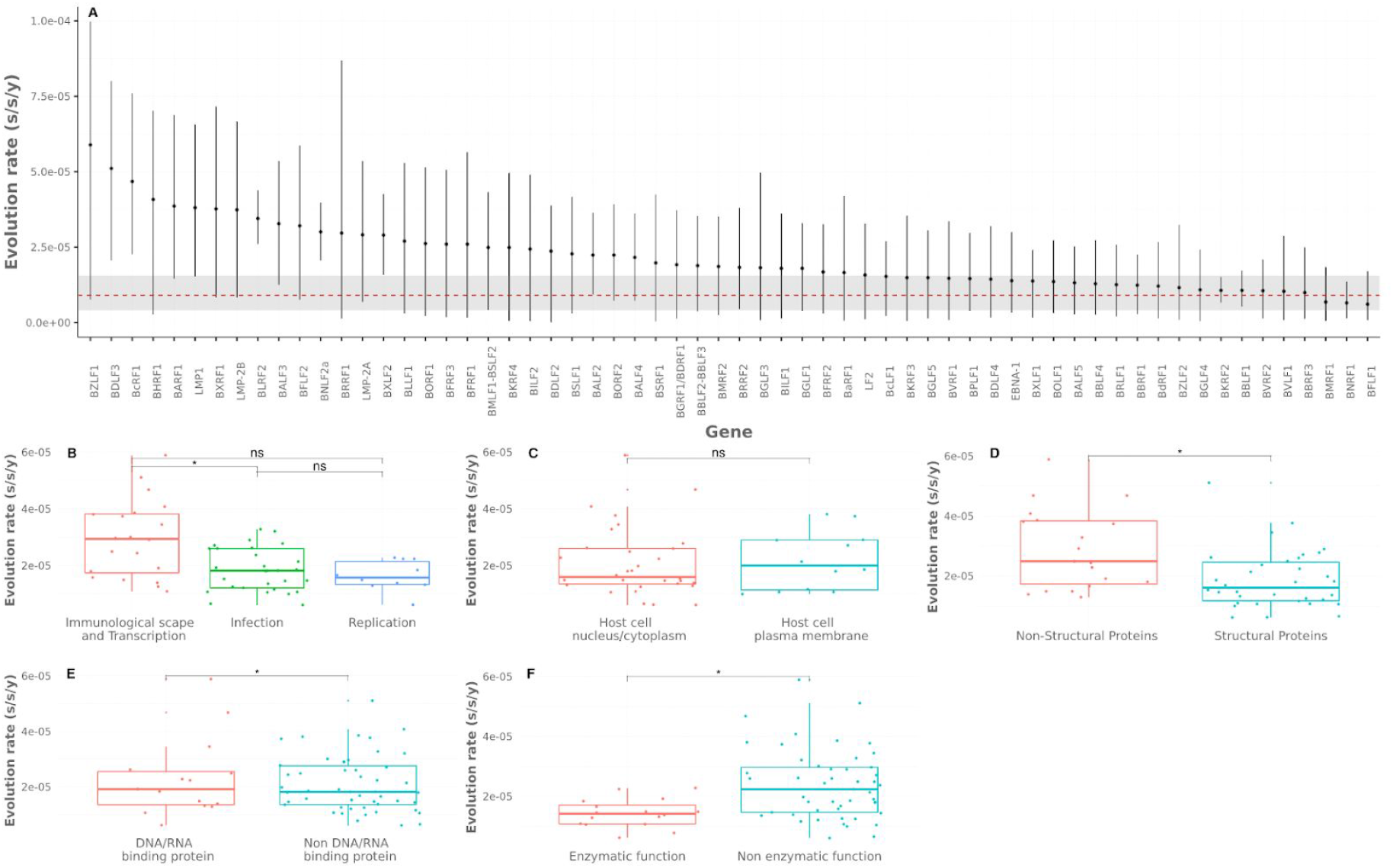
Evolutionary rates analysis. *(A)* Evolution rates for 60 analyzed genes. Dots depict mean evolution rates. Error bars indicate 95% HPDI (highest posterior density interval). Red dotted line depicts the genomes’ mean evolution rate, the gray horizontal stripe denotes the 95% HPDI for the genome’s mean evolution rate. *(B-F)* Comparison of gene’s evolution rates among different groups contained in the biological categories (gene names listed in table 1). Results are depicted in box plots. Asterisks denote statistical significance. Adjusted p-value □0.05 (*); adjusted p-value □ 0.01 (**); adjusted p-value >0.05 (ns).

For a comprehensive analysis of genes’ evolutionary rate, EBV genes were grouped based on the three genetic domains, as defined in GO database, by means of an unsupervised analysis. The groups were obtained by k-means method and refined and merged manually, considering their annotated function, in order to avoid redundancies and produce a better fit of the groups. In addition to those gene domains defined by the unsupervised analysis, two additional categories were defined on the basis of biological criteria, namely viral component and Enzymatic Function. The resulting groups are presented in Table 1.

**Table 1:** The table describes the groups as defined by the unsupervised analysis from the Gene Ontology database and the groups created based on their biological significance and those genes belonging to each group.

|  |
| --- |
| **Biological Process** |
| Viral immunological Scape and Transcription<br>LMP1, LMP2A, LMP2B, BGLF4, BZLF1, EBNA1, BMLF1-BSLF2, BILF1, BLRF2, BNLF2a, BGLF5, LF2, BCRF1, BHRF1, BRLF1, BcRF1, BARF1, BLDF3, BILF2, BRRF1 |
| Viral infective<br>BBRF3, BGLF3, BZLF2, BcLF1, BXLF2, BSRF1, BDLF1, BdRF1, BFLF1, BPLF1, BFRF3, BDLF2, BLLF1, BFRF1, BFLF2, BKRF2M BVRF1, BMRF2, BGLF1, BBRF2, BALF3, BGRF1/BDRF1, BALF4, BBRF1, BOLF1, BNRF1, BORF1, BVRF2, BBLF1 |
| Viral replicative<br>BMRF1, BSLF1, BBLF2, BALF2, BALF5, BXLF1, BaRF1, BORF2, BBLF4, BKRF3 |
| **Cellular component** |
| Host cell plasma membrane<br>BLLF1, BALF4, BXLF2, BKRF2, BMRF2, BBRF3, BBLF1, BZLF2, BILF1, LMP1, LMP2a, LMP2b |
| Host cell nucleus/cytoplasm<br>BMRF1, BGLF4M BPLF1, BFLF1, BLRF2, BXRF1, BGLF1, BBRF2, BSRF1, BDLF1, BORF1, BFRF3, BVRF1, BcLF1, BVRF2, BNRF1, BOLF1, BXLF1, BALF2M BBRF1, EBNA1, BZLF1, BHRF1, BcRF1, BSLFS/BMLF1, BALF5, BGLF5, BBLF4, BALF3, BKRF3, BSLF1, BaRF1 |
| Extracellular space<br>BCRF1, BARF1 |
| **Viral Component** |
| Structural proteins<br>BLLF1, BALF4, BXLF2, BKRF2, BMRF2, BBRF3, BBLF1, BZLF2, BDLF3, BDLF2, BFRF2, BKRF4, BdRF1, BGLF4, BMRF1, BPLF1, BFLF1, BLRF2, BXRF1, BGLF1, BBRF2, BSRF1, BDLF1, BORF1, BFRF3, BVRF1, BcLF1, BVRF2, BNRF1, BOLF1, BXLF1, BALF2, BBRF1, BALF5 |
| Non Structural proteins<br>LMP1, LMP2a, LMP2b, BILF1, BILF2, EBNA1, BZLF1, BcRF1, BCRF1, BARF1, BHRF1, BSLF2-BMLF1, BGLF5, BBLF4, BKRF3, BSLF1, BaRF1 |

| **Enzymatic Function** |
| --- |
| Enzymatic function<br>BGLF5, BGLF4, BGRF1/BDRF1, BPLF1, BKRF2, BVRF2, BSLF1, BORF2, BBLF2, BaRF1, BKRF3, BXLF1, BALF5, BBLF4, BLLF3, BMRF1 |
| Non enzymatic function<br>BZLF1, BDLF3, BcRF1, BHRF1, BARF1, LMP1, LMP2a, LMP2b, BLRF2, BNLF2a, BRRF1, BMLF1-BSLF2, BILF2, BGLF3, BILF1, BFRF2, BDLF4, EBNA1, BRLF1, BVLF1, BXRF1, BALF3, BFLF2, BXLF2, BDLF1, BLLF1, BORF1, BFRF1, BFRF3, BKRF4, BDLF2, BALF4, BSRF1, BMRF2, BBRF2, BGLF1, BcLF1, BVRF1, BOLF1, BBRF1, BdRF1, BZLF2, BBLF1, BBRF3, BNRF1, BFLF1, BALF2 |
| **DNA binding protein** |
| DNA/RNA binding protein<br>BGLF5, BGRF1/BDRF1, BKRF2, BSLF1, BBLF2M BALF5, BBLF4, BMRF1, BZLF1, BcRF1, BLRF1, BMLF1-BSRLF2, EBNA1, BORF1, BALF2 |
| Non DNA/RNA binding protein<br>BDLF3, BHRF1, BARF1, LMP1, LMP2a, LMP2b, BNLF2a, BRRF1, BILF2, BGLF3, BILF1, BFRF2, BDLF4, BRLF1, BVLF1, BXRF1, BALF3, BFLF2, BXLF2, BDLF1, BLLF1, BFRF1, BFRF3, BKRF4, BDLF2, BALF4, BSRF1, BMRF2, BBRF2, BGLF1, BcLF1, BVRF1, BOLF1, BBRF1, BdRF1, BZLF2, BBLF1, BBRF3, BNRF1, BFLF1, BGLF4, BPLF1, BVRF2, BORF2, BaRF1, BKRF3, BXLF1, BLLF3 |

#### Biological Process

three groups were obtained after the unsupervised analysis using GO Biological Process domain. The first group included genes involved in immunological escape. The second group contained genes implicated in the interaction between viral and host cells and the fusion of the viral and cell membranes. Finally, the third group includes only genes necessary for DNA viral replication.

#### Cellular Component

three groups were obtained performing the unsupervised analysis with GO Cellular component domain. The first group was defined by integral proteins located in the host cell membrane and/or in the virion membrane. The second group contained proteins that exert their action on the nucleus and cytoplasm of the host cell. The last group was defined by proteins present on the extracellular space; however this group was not further considered in the evolutionary rate analysis since it was composed by only two genes.

#### Viral Component

additionally two groups were defined by manual curation of GO Component Cellular domain: structural and non-structural.

#### DNA/RNA binding protein

only 31 genes were annotated with GO Molecular Function domain, by means of unsupervised analysis two groups were obtained, discriminated by the ability of gene products to bind DNA or RNA and gene products that don’t bind DNA or RNA.

#### Enzymatic Function

additionally two groups defined as enzymatic and non enzymatic functions were manually curated using GO Molecular Function.

The number of genes in the mentioned groups was further enlarged with genes with an annotated function in GenBank.

Using all the previously defined groups, the calculated evolutionary rate for each EBV gene was fitted into these categorical groups and the groups’ mean evolutionary rates were compared by pairs.

First we compared evolution rates among groups contained in the Biological Process category. Genes involved in viral immunological escape presented higher evolutionary rates than genes implicated in the entry and exit of the viral particle (Wilcoxon test, Bonferroni correction *P*=0.022, Fig. 6B) and also tends to be greater than genes associated with DNA viral replication (Wilcoxon test, Bonferroni correction *P*=0.074). No differences were found between mean evolutionary rates from genes needed for DNA biosynthesis and genes involved in the viral entry and viral assembly (Wilcoxon test, Bonferroni correction *P*=1).

Next we assessed differences in evolutionary rates of genes in the Cellular Component domain. Mutation rates of gene products located in cellular membranes didn’t differ from that of gene products localized in the cell nucleus (Wilcoxon test, Bonferroni correction *P*=1, Fig. 6C). Within the Viral Components group, those genes that code for non-structural proteins have a higher evolutionary rate than genes coding for structural proteins (Wilcoxon test, Bonferroni correction *P*=0.024, Fig. 6D).

When comparing evolutionary rates of genes in the DNA/RNA binding protein group, these evolutionary rates did not differ between gene products with the ability to bind DNA or RNA and those with no such ability (Wilcoxon test, Bonferroni correction *P*=1, Fig. 6E). Finally, the evolution rate for those genes with enzymatic function was smaller than that of those not coding for enzymatic proteins (Wilcoxon test, Bonferroni correction *P*=0.014, Fig. 6F).

These results, covering a gene-by-gene evolution rate calculus and its relation to their biological function, all together support the well-established notion that genes under constant immune pressure will have higher evolution rates in order to evade immune surveillance.

## Discussion

In this paper we sequenced 15 EBV complete genomes from pediatric patients from Argentina and analyzed their phylogenetic relation and variability pattern in the context of other complete genomes from different parts of the globe. Furthermore, we assessed the evolution rate of both, the entire genome and at a gene-by-gene level while exploring the biological significance of their evolution rate in relation to their functionality or localization of the gene product to the cell and/or viral particle.

Other research groups have previously generated up to 1000 complete EBV genome sequences from different geographic locations (Correia et al. 2018; Wegner et al. 2019); however, each group analyzed their datasets in an independent way without any standardized criteria. This fact represents no problem for each individual analysis, but becomes a major concern when integrating all independently generated consensus sequences for further analysis. There is no unique, gold-standard, bioinformatic pipeline to obtain the final EBV genome consensus sequence, something of particular importance given the amount of repetitive regions in EBV genome, and that different research groups would obtain different consensus genomes if analysing the same dataset. For instance, systematic differences were observed in the way that each group process repetitive regions of the genome, deletions sites or poorly covered regions, where some bioinformatic strategies replace the lack of a good quality sequence by the reference sequence, gaps or with unknown nucleotides (N) or by cutting off that region, all of which will have a different meaning in a post phylogenetic analysis. This discrepancy becomes most relevant when comparing sequences from different sources as we did in this study. In order to produce higher quality results we have demonstrated the importance of analysing the raw fastq data files using one same pipeline instead of aligning the consensus sequences directly downloaded from NCBI. In this scenario, our new pipeline took special consideration of these issues by masking conflictive regions with N nucleotides and conserving small insertions and deletions (Indels), since it retained this information to produce the final consensus genome.

The sequencing of complete EBV genomes from Argentina and their integrated phylogenetic analysis with genomes from other regions of the world revealed the presence of two groups supported by a high bootstrap value, a small group formed by EBV2 sequences and a much larger group composed by EBV1 sequences. From the full data set, 18 sequences were from Argentina, 15 newly sequenced and 3 downloaded from NCBI. These sequences were found in both EBV1 and EBV2 groups and were interspersed among sequences from different geographic locations, indicating their independent origins. This work contributes to enlarge the amount of fully characterized EBV genomes from our region, especially from Argentina, which were until now underrepresented in the NCBI database; moreover, reporting new EBV2 genomes is of particular highlight since EBV2 from South America are even scarcer (Wegner et al. 2019; Correia et al. 2018).

After restricting the analysis to EBV1 genomes, a geographical structure arose for this viral type from the phylogenetic and principal component analysis. In the former case, bootstrap supports allowed for the discrimination of the EBV1 clade into a grate Asian clade, which could be further differentiated into four smaller sub-clades (Western Asia, Eastern Asia, Southeast Asia and Europe/Australia). On the other hand, a cosmopolitan clade contained genomes from Africa, Europe, Australia and the Americas. In the latter case, the PC1 mainly separated the Asian sequences for the rest of the geographies, but was able to further discriminate the Asian group into a Southeast Asia group and another group composed of Eastern and Western Asian genomes. Furthermore, the PC2 could further discriminate Southeast Asian sequences from the rest of Asia. This geographical structure is in accordance with previous observations (Bridges et al. 2019), (Wegner et al. 2019), even after the inclusion of our 15 Argentinean sequences, which denotes a high segregation potential of the Asian sequences; another interpretation of this fact could be a high resemblance of American sequences with those from Africa, Australia and Europe. This latter interpretation could be plausible given the imperialist history of humanity and historical movements of human, since the American, African and the Australian continents suffered large-scale population changes from European colonialism and migrations (Gilmartin, n.d.). In particular, an estimated 1.5 million European migrants arrived to the Americas along with the introduction of an estimated 7 million African slaves between XV and XVIII centuries (Avena et al. 2012). It is worth mentioning that having possible recombinant sequences removed from the alignment strengthens our observations, since recombinant genomes could distort the topology of the tree, as demonstrated by Zanella et. al. (Zanella et al. 2019).

The fact that Asian sequences presented higher variability than those from other regions, with respect to the EBV wild-type reference, and that those variants were distributed along the entire genome, rather than in a single gene, could support the notion of demographic events during ancestral human migrations. In addition, the low internal diversity observed among Asian sequences might suggest the existence of a bottleneck event. Remarkably this evolutionary pattern was also observed for other herpesviruses, namely HSV1, HHV8 (Liu et al. 2017; Kolb, Ané, and Brandt 2013) as well as for other viruses (Firth et al. 2010).

Altogether, these results support the idea that one EBV reference genome does not fully represent wild type EBV from all the geographies.

Previously, using first-generation gene sequencing techniques some latent genes have been shown to contribute to the differential segregation of Asiatic sequences (Chang et al. 2009; Tzellos and Farrell 2012). Nowadays, massive sequencing techniques allow us to identify variants that prevail in each geography and their distribution in a whole genomic scale, enabling us to detect those genes mainly contributing to this clustering. Given the prior observations in the phylogenetic tree and PCA analysis, a DACP analysis was implemented to disclose which genomic positions contribute to the segregation of the two main groups, the Asian and the cosmopolitan non-Asian clades. To the best of our knowledge, this is the first time that the contribution of each single gene or group of genes was assessed in the context of the entire EBV1 genome in this way. The analysis revealed four main genomic regions with geographic segregation power. Variants in the first genomic region were mainly concentrated in the BPLF1 gene. Previous work has reported a large content of non-synonymous variants in this gene when assessing gastric carcinoma and control samples, but no association between them was disclosed until now (Yao et al. 2017). Nonetheless, the authors suggest that BPLF1 gene could be under positive selection, although the variability of this gene in the geographical context had not yet been reported. Variants present in the second group were identified to the terminal region of the EBNA-1 gene. The EBNA-1 gene has been thoroughly sequenced in order to study its association with malignant pathologies and geographies and has proven to be a good molecular marker of geographic origin, based on aminoacid position 487 in the C-terminal region of the protein (Correia et al. 2017; Mario Alejandro Lorenzetti et al. 2010; Chang et al. 2009). Our present results are in accordance with these previous observations that EBNA1 gene is a strong geographical indicator.

Concerning the region located between positions 120,000 and 140,000 of the reference genome, the main geographical segregating SNPs were located in the BDLF3, BcRF1 and BXRF1 genes, mainly late phase genes; however, as far as we know, there are no reports concerning these genes variation with respect to their geographic origin.

Finally, the last region with geographic segregation potential mapped to the latent membrane proteins, LMP1, LMP2A and LMP2B. Remarkably, the geographical variability of these genes has been previously well characterized in different genetic approaches, not considering the weight of each individual genes’ segregation contribution, as influenced by the other genes in the genome (Correia et al. 2017; Chang et al. 2009). Moreover, our group previously characterized a distinct LMP1 variant of preferentially circulating in our country (Gantuz et al. 2017), a fact that highlights the actual value of enlarging the proportion of complete genomes from our country, in order to overcome the current scarcity of data from South America.

In this work, the evolution rate of each gene and for the entire genome were calculated in order to characterize those genes with higher mutational rates. Given that the EBV genome is made up of a large double-stranded DNA molecule (dsDNA), approximately 172 kb, and has a DNA polymerase with a reliable proof-reading activity, a high degree of conservation was expected, as previously demonstrated (Lynch 2010; Sanjuán and Domingo-Calap 2016). This analysis requires a large number of sequences covering an ample scale of time, in order to achieve an appropriate temporal structure for phylodynamic analysis. From the original data set, only 55 sequences had information on the isolation date, this being the main limitation to understanding the evolution over time of each of the EBV genes. There are two possible strategies to overcome this limitation: one of them is to assume the co-evolution of the EBV genome and the host genome. The second possible strategy is to calibrate the analysis with an evolution rate reported in the bibliography for a particular gene of the same virus or of an evolutionarily close virus, estimating the age of the common ancestor and then calibrating the remaining analyzes with these data (Lee and Ho 2016). In particular, our analysis was performed using the second strategy, because with the assumption of a virus-host co-evolution, the viral evolutionary rate would be forced to be on the same evolutionary scale as that of its hosts. On the contrary, the second analysis strategy makes it possible to independently estimate the viral evolutionary rate. In our study, which made use of 55 EBV isolates from different geographic regions, we estimated a substitution rate for the entire EBV genome (9.09e-6 s/s/y), two orders of magnitude higher than that of the human host (0.5e-9 s/s/y) (Scally 2016), and similar to that of other double-stranded DNA viruses, including herpesviruses (Firth et al. 2010). This result suggests that EBV co-evolving together with humans is not necessarily the most appropriate assumption for this virus, in accordance to a previous observation for HSV-1, VZV (Firth et al. 2010).

In addition, the evolutionary rate for most individual genes was evaluated, being BZLF1, BcRF1, BDLF3, BXRF1 and LMP1 among those that showed the highest evolution rates. Notably, these genes were also the same ones most contributing to both, viral type segregation (after EBNA2 and EBNA3s) and geographic segregation.

Last, but not least, we analyzed the evolutionary rates in terms of functional gene groups, which were constructed according to the GO domains. Not surprisingly, our analysis showed that genes coding for viral immunoescape-related proteins have accumulated a higher number of variants over time; a selected characteristic that may have provided the virus with the ability to latently persist for a lifetime in the infected host. As expected, genes coding for enzymes showed lower evolutionary rates, which translates into a high level of conservation given the necessity for functional enzymes in order for EBV to undertake sporadic reactivation and replication cycles. In addition, it is worth mentioning that genes coding for structural proteins (proteins packed in the viral particle) showed evolutionary rates significantly lower than non-structural ones, another characteristic of out-most evolutionary importance given that structural proteins will determine the success of early infection stages. Moreover, this fact was consistent with the previous observation, since most gene products involved in immunoescape are indeed non-structural proteins.

Given that circulating EBV genomes in South America are underrepresented, this work constitutes a valuable contribution to a better understanding of the relationship between viral variability, geography related demographic events. Also, a large number of EBV2 sequences have been contributed, but even so the number of sequenced genomes of this viral type continues to be small, making statistical analyses not plausible and therefore not possible to reach a reliable conclusion. Past events such as the arrival of the European to America and more recent events such as human migrations could explain the relationships observed in phylogenetic analysis.

In short, this work contributed to overcome the scarcity of EBV complete genomes from South America, and hence, achieved the most comprehensive geography-related variability study by determining the actual contribution of each EBV gene to the geographic segregation of the entire genomes. Moreover, to the best of our knowledge, we established for the first time the evolution rate for the entire EBV genome and statistically demonstrate that evolution rates, on a gene-by-gene basis, are related to the encoded protein function. Considering evolution of dsDNA viruses with a codivergence-independent approach, may lay the basis for future research on EBV evolution. Last but not least, this work also expands the sampling-time lapse of available complete genomes derived from different EBV-related conditions, a matter that until today, prevents for detailed phylogeographic analysis.

## Methods

### Ethics statement

Hospital’s ethics committee reviewed and approved this study (*CEI Nº 18.04*), which is in accordance with the human experimentation guidelines of our institution and with the Helsinki Declaration of 1975, as revised in 1983. Clinical samples from patients with EBV infection were anonymized prior to this study. A written informed consent was obtained from all the patient’s parents or tutors.

### Patients and samples

This study included samples from 15 pediatric patients with EBV related diseases from Argentina: i. three children with infectious mononucleosis (IM) with a median age of 4 years (range, 1 to 17 years) and 52% male; ii. a child with secondary immunodeficiency following liver transplant; iii. eleven patients with EBV-positive lymphomas (6 Hodgkin and 5 non-Hodgkin) with a median age of 9 years (range, 3 to 18 years), and 77% males.

The IM cases were identified on clinical grounds and confirmed by indirect immunofluorescent assay (IFA) for the detection of IgM antibodies against virus capsid antigen (VCA) on commercial EBV-VCA antigen substrate slides (MBL BION, EB-5012) according to the manufacturer’s instructions. The liver transplant patient was followed-up with a periodic monitoring of viral load at our institution. Six ml of EDTA-peripheral blood and a pharyngeal secretion sample were obtained from patients with IM and from the liver transplanted child.

The association between the lymphomas and EBV was examined on a previously formalin-fixed and paraffin-embedded tissue sections of the same tumor biopsy using the PNA *in situ* hybridization (ISH) kit (Dako, K5201) together with a PNA-probe to detect Epstein-Barr encoded RNAs (EBERs) (Dako, Y5200), according to the manufacturer’s instructions. Cases were rendered as EBV-associated when specific staining was observed in the nucleus of tumoral cells without staining in infiltrating lymphocytes. In addition, fresh frozen tumor tissue biopsies were obtained from our institutional tumor repository.

### DNA Extraction

Total DNA was purified from Ficoll-isolated PBMCs, pharyngeal secretions and fresh frozen tumor biopsies with the QIAamp DNA minikit (QIAGEN, 51304), according to the manufacturer’s instructions.

### Viral load assays

EBV genome copy number was measured in all samples by a real time quantitative PCR (qPCR) assay, as previously described (Fellner et al. 2016) on a LightCycler 480 device. Standard curve was performed with 1/10 serial dilutions of an EBNA1-pGEM-T Easy plasmid, a kind gift from Dr. Dolores Fellner, and ranged from 10^7^ to 10^2^ EBNA1 gene copies. Finally, viral load was calculated as EBV genome copy number per μg of total genomic DNA. Samples with a viral load greater than 10^6^ copies/ug DNA were selected for library preparation and sequencing. In those cases where PBMCs and pharyngeal secretions were available, we selected the sample with a higher viral load assuming neglectable viral variation between both anatomical compartments, as previously observed (M. A. Lorenzetti et al. 2014).

### Library preparation and Illumina sequencing

Sequencing libraries were constructed with SureSelect QXT Target Enrichment kit (Agilent Technologies, G9683B) and custom designed EBV RNA-bait probes (Agilent Technologies, 5190-4806), according to the manufacturer’s instructions. RNA-bait probes were designed and kindly shared by Dr. Daniel P. Depledge (Depledge et al. 2011). EBV-enriched libraries were sequenced on a NexSeq500 Illumina device, with 300 cycle mid-output (2×150 bp) NextSeq Reagent kits v2 (Illumina Corporation, 20024905). FASTQ files of raw sequencing data were deposited in NCBI database with the Sequence Read Archive (SRA) accession number PRJNA679281 (Supplemental Table 4).

### Public available sequencing data

Publicly available raw NGS data were downloaded from SRA-NCBI database (Supplemental Table 4). Downloaded data belonged to BioProjects: PRJNA522388, PRJNA505149 and PRJEB2768.

### Bioinformatic data processing

The bioinformatic analysis was implemented with a mapping-based customized pipeline designed at the bioinformatics core of our institution. Initial preprocessing steps, including read quality checks and sequence trimming, were performed with fastp software v.0.20.1 (Chen et al. 2018). Duplicate removal was assessed with PRINSEQ software v.0.20.4 (Schmieder and Edwards 2011). Thereafter, reads were mapped to both EBV-type reference genomes (NC_007605.1 for EBV1 and DQ279927.1 for EBV2) making use of Burrows-Wheeler Aligner (BWA) v.0.7.17-r1188 (Li and Durbin 2009) which generates BAM format output files. After that, BAM files were processed using SAMTools v1.9 (Li et al. 2009). This phase utilizes both available reference genomes and reports the number of variants over EBNA2 and the entire EBNA3 family genes to establish EBV type. Variant calling step and consensus sequence generation were performed with bcftools mileup samtools software package v.1.7 (Li et al. 2009). In addition, our pipeline takes into consideration masked repetitive regions, as well as the uncovered ones to avoid biases in the detection procedure of indels.

After a mapping step against both reference genomes, EBV type was predicted based on the number of variants considering EBNA2 and the entire EBNA3 family of genes. This typing step uses the VCF file and operates under the assumption that a greater number of genetic variants will arise when mapping against the wrong type reference; on the contrary, a better mapping quality will result when mapping against the correct reference (Supplemental Table 5).

Novel NGS data from samples sequenced in this study, as well as raw data downloaded from NCBI were analyzed with the same bioinformatic pipeline.

### Multiple sequence alignment and evolutionary tests

Multiple sequence alignment (MSA) was created with MAFFT v7.310 (Katoh and Standley 2013) and cured manually for further gap reductions.

Aligned genomes were assessed for potential recombination signals with RDP4.101 software (Martin et al. 2015), and those sequences deemed as potentially recombine/ant genomes were removed from the alignment.

IQ-TREE software v1.6.1 (Nguyen et al. 2015) was used to determine the best partition schemes for the MSA and to predict the best evolutionary model for each partition (Supplemental Table 6). Then phylogenetic reconstruction was performed under the maximum likelihood (ML) method and 1000 ultrafast bootstrap iterations were performed. The ML ultrafast bootstrap tree was visualized in FigTree v1.4.3 (http://tree.bio.ed.ac.uk/software/figtree/). Geography-based segregation analysis of EBV genomes was performed with iTOL (Letunic and Bork 2016).

The alignment was imported into R software v4.0.2. (R Development Core Team 2003) using the ape package and the SNPs were extracted using the adegenet package (Jombart 2008). Principal components analysis (PCA) and Principal Component Discriminant Analysis (DAPC) were performed with the adegenet package.

The EBV1 reference sequence was compared with sequences belonging to the same geographical region with Consensus Compare tool, kindly provided by Dr. PJ Farrell, to evaluate specific geographical polymorphisms.

### Estimation of evolution rate

Fifty five sequences with sampling date, ranging from 1963 to 2019, and from different regions of the world were further selected for evolution rate estimation (10 EBV2 and 45 EBV1). Since this new dataset presented no temporal structure, a previously published evolution rate for LMP1 gene-obtained from data with temporal structure-was used for molecular clock calibration (Gantuz et al. 2017). Time to the most recent common ancestor (tMRCA) for LMP1 was initially estimated and used to calibrate the rest of the analyses. Evolutionary models were estimated with IQ-TREE software v1.6.1 (Supplemental Table 7). Bayesian coalescent analysis was implemented in BEAST v1.8.4 (Drummond et al. 2012) using an uncorrelated lognormal relaxed clock model (UCLN) and a Gaussian Markov random field (GMRF) Bayesian Skyride demographic model. Analyses were run for 80 and 100 million iterations of Markov Chain Monte Carlo (MCMC). Results were visualized in Tracer v-1.7 (Rambaut et al. 2018) and convergence was assessed with effective sample sizes higher than 200.

### Gene Ontology-based clustering

Genes were grouped using an unsupervised procedure that took into consideration three well distinguished Gene Ontology (GO) domains: Biological Process (BP), Cellular Component (CC) and Molecular Function (MF). First, direct EBV gene annotations were obtained from the Uniprot database (https://www.uniprot.org/) and extended to all its parent relationships among each domain. This information was summarized by means of a binary matrix constructed for each GO domain. Then, a gene to gene distance matrix for each domain was computed based on the Jaccard index. Over these matrices, the presence of non random structure was explored using both, the Hopkins statics and Visual Assessment of cluster Tendency (VAT), in clustertend v1.4 (Lawson and Jurs 1990) and factoextra v.1.0.7 (Kassambara 2017) R packages. Finally, a K-means algorithm was used to undertake gene groups for each GO domain. The optimal number of partitions (K) was initially explored through three quantitative methods (Elow, gap statics and average silhouette) and then refined manually by merging clusters with redundant biological interpretation.

### Statistical analysis

Statistical analysis was performed using the computing environment R version 4.0.2. Normality and homoscedasticity were evaluated using Shapiro-Wilk and Bartlestt test, respectively. Groups that meet these principles were analyzed with the t-student test and groups that didn’t meet these criteria were compared with Wilcoxon, or Kruskal-Wallis tests, as appropriate. All *P*-values were adjusted using the Bonferroni test.

## Acknowledgment

This study was supported in part by a grant from the National Agency for Science and Technology Promotion (ANPCyT) (PICT 2016 N°0548). A.J.B, M.A.L, and M.V.P are members of the CONICET Research Career Program. A.C.B. is a fellow from ANPCyT.

We extend a very special thanks to Dr. Judith Breuer and all members of the Pathogen Genomics Unit at University College London, for receiving M.A.L in the lab and the training in NGS library preparation with target enrichment techniques and NGS sequencing of EBV.

This training was supported by the Jorge Oster grant from “Fundación Bunge y Born”.

